# The role of environmental calcium in the extreme acid tolerance of northern banjo frog (*Limnodynastes terraereginae*) larvae

**DOI:** 10.1101/2022.04.18.488693

**Authors:** Coen Hird, Craig E. Franklin, Rebecca L. Cramp

## Abstract

Many aquatically respiring animals inhabiting low pH waters can suffer acute inhibition of ion uptake and loss of branchial (gill) epithelial integrity, culminating in a fatal, rapid loss of body Na^+^. Environmental calcium levels ([Ca^2+^]_e_) are pivotal in maintaining branchial junction integrity, with supplemental Ca^2+^ reversing the negative effects of low pH in some animals. Tolerance of some naturally acidic environments by aquatic animals is further complicated by low [Ca^2+^]_e_, yet many of these environments are surprisingly biodiverse. How these animals overcome the combined damaging actions of low pH and low environmental Ca^2+^ remains unknown. Here, we examined the effects of [Ca^2+^]_e_ on the response to low pH in larvae of the highly acid tolerant frog *Limnodynastes terraereginae*. Acute exposure to low pH water in the presence of low [Ca^2+^]_e_ increased net Na^+^ efflux. Provision of additional [Ca^2+^]_e_ reduced net Na^+^ efflux, but the effect was saturable. Acclimation to both low and high [Ca^2+^]_e_ improved the resistance of larvae to Na^+^ efflux at low pH. Inhibition of apical Ca^2+^ uptake by ruthenium red resulted in an abrupt loss of tolerance to low pH in larvae acclimated to low pH water. Acclimation to acidic water increased branchial gene expression of the intracellular Ca^2+^ transport protein calbindin, consistent with a role for increased transcellular Ca^2+^ trafficking in the tolerance of acidic water. This study confirmed the physiological challenge of low [Ca^2+^]_e_ on branchial integrity in acidic waters and highlighted a potential role for maintenance of transcellular Ca^2+^ uptake in the acid tolerance of *L. terraereginae*.

**Summary statement:** Tolerance of naturally acidic, dilute, and soft waters by larvae of the frog *Limnodynastes terraereginae* involves adaptations to the branchial calcium transport pathway which protects intercellular junctions against damage.

## INTRODUCTION

Life in freshwater environments is complicated by the fact that animals are hyper-ionic with respect to the environment and ions tend to move out of the animal along their diffusion gradients. In order to offset ion losses, freshwater animals actively take up ions from the environment via specialised cells in the gills, integument (in larvae and embryos) and across the gut (Edwards and Marshall, 2012; Evans et al., 1999). The physiological challenges of living in freshwater environments can be further compounded by low pH. Most aquatically respiring animals are intolerant of water pH less than 5 (low pH). In many animals low pH water substantially impairs both ion uptake capacity and epithelial integrity, leading to a rapid loss of homeostasis that can be fatal (Freda and Dunson, 1984; McDonald et al., 1984; Meyer et al., 2010; Meyer et al., 2020; Robinson, 1993).

The effect of low pH water on branchial epithelial integrity drives its toxicity in many species. In acid-sensitive animals, exposure to low pH water reduces transepithelial resistance through the disruption of paracellular tight junctions (Daye and Garside, 1976; Meyer et al., 2010; Rosseland and Staurnes, 1994). Tight junctions (TJ) are composed of a complex of transmembrane and associated proteins which function in cell-cell adhesion and control epithelial permeability by acting as a selective barrier to ions and small molecules (Schneeberger and Lynch, 2004). In doing so, TJs regulate and maintain key transcellular ion gradients necessary to facilitate transepithelial ion uptake. Low extracellular pH has been shown to reduce the resistance of the paracellular pathway by changing the conformation of TJ proteins which alters their gating properties (Schneeberger and Lynch, 1992; Schneeberger and Lynch, 2004).

Environmental [Ca^2+^] is essential to maintaining the integrity of the epithelial TJ (Schneeberger and Lynch, 1992). Increased [Ca^2+^]_e_ has been shown to ameliorate the effects of low pH on epithelial integrity in some fish and amphibian species (Dalziel et al., 1986; Matsuo and Val, 2002; McDonald et al., 1983; Meyer et al., 2020). This may be because low [Ca^2+^]_e_ causes dissociation of the paracellular junctions, increasing epithelial permeability (Bhat et al., 1993; Ma et al., 2000; O’Keefe et al., 1987). In some fishes and larval amphibians acutely exposed to low pH, elevated and reduced environmental Ca^2+^ levels substantially reduced and increased rates of passive net Na^+^ loss, respectively (Cummins, 1988; Freda et al., 1991; Gascon et al., 1987; Gonzalez and Dunson, 1989; Kumai et al., 2011; McDonald and Rogano, 1986; McDonald et al., 1983). Ca^2+^ may act on the TJ directly, but also indirectly through Ca^2+^-dependent proteins in adherens junctions (AJ) which sit basally to the TJ (Brown and Davis, 2002). Removal of Ca^2+^ from the extracellular space has been shown to reduce cytosolic [Ca^2+^] (Gonzalez-Mariscal et al., 1990), cause the detachment and internalisation of extracellular AJ and TJ proteins (Volberg et al., 1986), and reduce transepithelial resistance (Gonzalez-Mariscal et al., 1990). Adherens junctions function primarily in cell-cell adhesion, but also the regulation of the actin cytoskeleton and as transcriptional regulators (Hartsock and Nelson, 2008). Extracellular Ca^2+^ can regulate paracellular permeability by interacting directly with calcium-dependent junctional proteins on the cell surface and/or through active transcellular Ca^2+^ uptake and cytosolic Ca^2+^ signalling pathways (Stuart et al., 1996). Although extracellular [Ca^2+^] is important in determining the effects of low pH exposure in fish and amphibians, the specific mechanism through which this occurs remains unknown.

Transcellular uptake of Ca^2+^ from the surrounding water via the gills constitutes the major route by which Ca^2+^ is taken up in fish (Baldwin and Bentley, 1980; Chasiotis et al., 2012; Flik and Verbost, 1995). Active branchial Ca^2+^ uptake occurs at ionocytes in the branchial epithelium (Fig. 1). Extracellular Ca^2+^ enters ionocytes through non-voltage gated epithelial Ca^2+^ channels (ECaC) in the apical membrane (Edwards and Marshall, 2012; Flik and Verbost, 1995) and is then shuttled to the basolateral membrane by Ca^2+^ binding proteins such as calbindin for extrusion via Na^+^-Ca^2+^ exchangers (NCX) and plasma membrane Ca^2+^ ATPase transporters (PMCA). Once in the basolateral extracellular space, Ca^2+^ can directly interact with the extracellular Ca^2+^ binding domain of E-cadherin in AJs to influence its properties and those of the overlying TJ (Pokutta et al., 1994; Zhang et al., 2009). Branchial Ca^2+^ absorption is tightly controlled through the regulation of ECaC activity and changes in the abundance of intracellular Ca^2+^ transport proteins (Cai et al., 1993; Kelly and Wood, 2008; Khanal and Nemere, 2008; Shahsavarani and Perry, 2006; Verbost et al., 1993; Wongdee and Charoenphandhu, 2013). This, in turn, may reduce the transcellular movement of Ca^2+^ to the basolateral extracellular space, which could limit its availability to E-cadherin in the AJ and compromise the permeability of the overlying TJ. Factors that compromise the maintenance of transcellular Ca^2+^ transport pathways could influence the Ca^2+^-sensitive aspects of junctional stability.

**Fig. 1.**
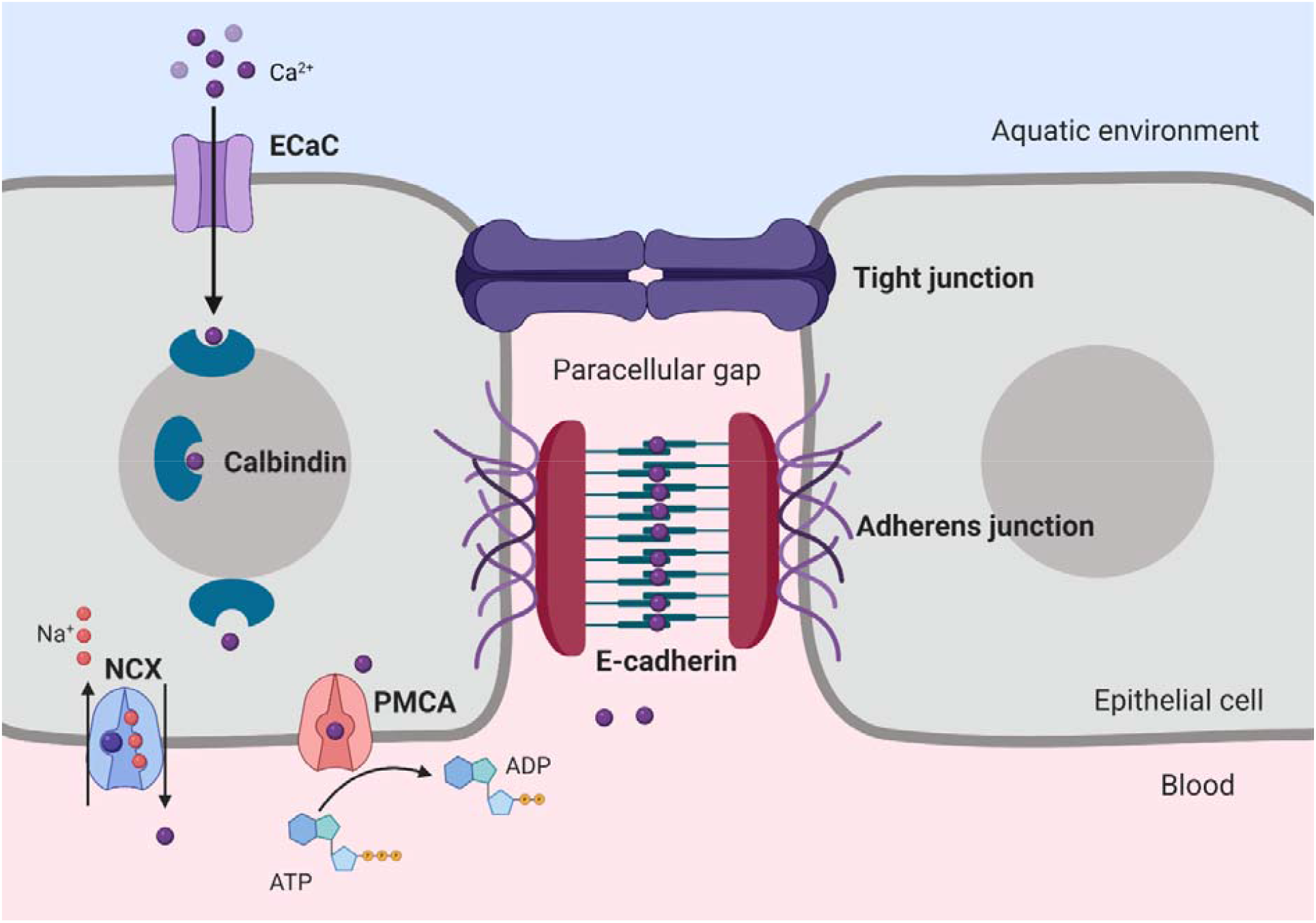
Proposed model of epithelial Ca^2+^ uptake and cell-cell adhesion in freshwater amphibian larvae ionocytes. Ca^2+^ is absorbed through the apical Ca channel (ECaC) and extrusion through basolateral Ca^2+^ transporters (NCX and PMCA) is facilitated and buffered by intracellular Ca^2+^ binding proteins (such as calbindin). Epithelial ionocytes are bound by the junctional complex consisting of tight junctions, and more basal adherens junctions which are Ca^2+^-dependent and regulate tight junction structure and function.

Given that highly acidic water is toxic to most aquatic animals, naturally acidic freshwater bodies are surprisingly biodiverse. Among the most acidic freshwater ecosystems in the world is the Wallum along the eastern coast of southern Queensland and northern New South Wales in Australia. Wallum ecosystems are characterised by highly acidic waters, ranging from pH 2.8 – 5.5 (Hines and Meyer, 2011). Compounding the difficulties of living at low pH, Wallum waters are also dilute (low in salts) and soft (low in Ca^2+^ and Mg^2+^; Bayly, 1964). Despite these challenges, larvae of some Wallum frog species can tolerate exceptionally acidic waters (Hines and Meyer, 2011; Meyer, 2004; Meyer et al., 2020). One such species is the northern banjo frog, *Limnodynastes terraereginae*, populations of which can be found throughout eastern Australia inhabiting aquatic environments which range in pH from circum-neutral to pH 3.0, making it one of the most highly acid tolerant vertebrate species known. Given that [Ca^2+^]_e_ is limited in Wallum environments and that Ca^2+^ uptake is typically inhibited by low pH, understanding how amphibian larvae manage Ca^2+^ transport in low pH water is likely central to understanding the mechanistic basis of their tolerance to these extreme environments.

To determine the importance of [Ca^2+^] on Na^+^ homeostasis at low pH, we examined whole animal Na^+^ and Ca^2+^ fluxes following both acute and chronic exposure to low pH and different [Ca^2+^]_e_ in *L. terraereginae* larvae. To understand the role of transcellular Ca^2+^ uptake for the maintenance of epithelial integrity at low pH, *L. terraereginae* larvae reared at low pH were exposed to the calcium channel antagonist, ruthenium red (RR). We also measured gene expression patterns of four key Ca^2+^ transport proteins (ECaC, calbindin, NCX, and PMCA) and E-cadherin in the gill epithelia. We hypothesised that acute exposure to low pH would result in increased net Na^+^ efflux under low [Ca^2+^]_e_, but that chronic exposure (acclimation) to low [Ca^2+^] would reduce net Na^+^ efflux with low pH exposure and increase Ca^2+^ influx. In addition, we hypothesised that the acute impairment of apical Ca^2+^ uptake via ECaC (with RR) would increase net efflux of Na^+^ at low pH consistent with a role for transcellular Ca^2+^ transport in the maintenance of intercellular junction integrity. Finally, acclimation to both low pH and low [Ca^2+^]_e_ was hypothesised to correspond to an increase in the expression of the four key Ca^2+^ transporters (ECaC, calbindin, NCX, and PMCA) and E-cadherin consistent with an increased rate of transcellular Ca^2+^ uptake and reinforcement of Ca^2+^-dependent AJs to protect junctional integrity in acid-acclimated larvae.

## MATERIALS AND METHODS

### Experimental animals and general methods

All animals were collected under the Queensland Department of Environment and Heritage Protection Scientific Purposes Permit (WITK15563515), and all procedures were approved by The University of Queensland’s Animal Ethics Welfare Unit (SBS/484/17). Limnodynastes terraereginae egg masses were collected in January 2018 from Bribie Island National Park, Queensland. Eggs were allocated to circumneutral (pH 6.5) or low (pH 3.5) pH artificial soft water (ASW; Freda 1984); distilled water plus (in μmol L^-1^) 40 CaCl_2_ .2H_2_O, 40 MgSO_4_.7H_2_O, 120 NaCl, 50 NaOH, 20 KCl; pH was adjusted with 0.1 M H_2_SO_4_). Hatched tadpoles were housed in 5 L plastic tanks connected to two 200 L filtered recirculating aquarium systems (15 tanks per system). Each system was connected to a canister filter for biological, mechanical, and chemical filtration (Fluval G6, City, Country). System pH was monitored daily (LAQUA P-22, Horiba Instruments, Singapore) and regulated as necessary through the addition of 0.1 M H_2_SO_4_. Water [Na^+^] and [Ca^2+^] was measured weekly using flame photometry (BWB Technologies, Berkshire, UK). Tadpoles were fed every second day with thawed frozen spinach, and each system underwent a 20% water change weekly. Room temperature was maintained at 22 ± 1°C with fluorescent overhead lighting programmed to a 12L:12D photoperiod (6:00 – 18:00). The following experiments were replicated once within the laboratory.

### Whole animal Na^+^ and Ca^2+^ fluxes

The Na^+^ and Ca^2+^ concentration (μmol L^-1^) in water samples was measured using a flame photometer (BWB Technologies, Berkshire, UK). Na^+^ and Ca^2+^ detection ranges were calibrated using (in μmol L^-1^) 5, 50, 500, and 1000 standards. To buffer potential spectral interference of sulphites and phosphates on Ca^2+^ readings, 50 μmol L^-1^ [Ca^2+^] samples were diluted to 50% and 250 μmol L^-1^ [Ca^2+^] samples were diluted to 25% (Thiers and Hviid, 1962; Welch et al., 1990). Water samples containing 5 μmol L^-1^ [Ca^2+^]_e_ were under the limit of detectability for our flame photometer and Ca^2+^ fluxes were unable to be reliably measured in this treatment; Na^+^ concentrations remained within the limits of detectability for all [Ca^2+^]_e_ exposures and were recorded for all treatments. All net Na^+^ and Ca^2+^ flux measures (in nmol L^-1^ h^-1^) were calculated by comparing changes in water [Na^+^] and [Ca^2+^] over the exposure period as follows:

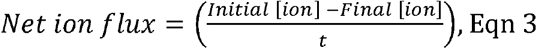

where *t* (in h^-1^) is the time of the exposure period in hours (6).

### Effects of acute exposure to high or low Ca^2+^ levels on whole animal net Na^+^ and Ca^2+^ fluxes at low pH (experiment 1)

To assess the effect of acute exposure to high or low [Ca^2+^]_e_ on acid-induced net Na^+^ and Ca^2+^ fluxes, *L. terraereginae* larvae (n = 36; Gosner stages 26-38; Gosner, 1960) were randomly allocated to three 5 L tanks (n ≤ 12 per tank to obtain adequate statistical power, based off previous ion flux work with the species in Meyer et al. 2004) containing 50 μmol L^-1^ [Ca^2+^]_e_ ASW (Table 1). Water pH was maintained at pH 6.5 and larvae were acclimated to these conditions for 4 weeks. Water pH was monitored daily and regulated as necessary via addition of 0.1 M H_2_SO_4_. Tadpoles were fed every second day with thawed frozen spinach, and each system underwent a 100% water change weekly. Ammonia levels were monitored using an API® Ammonia Test Kit (Mars Fishcare, Chalfont, PA). Larvae were then placed into individual 200 ml glass beakers containing 50 ml of either 5 μmol L^-1^ [Ca^2+^]_e_, 50 μmol L^-1^ [Ca^2+^]_e_ or 250 μmol L^-1^ [Ca^2+^]_e_ ASW (n = 12 per treatment) 30 minutes prior to testing. Water pH in half of the beakers in each treatment (n = 6) was acutely lowered to pH 3.5 through the addition of dilute H_2_SO_4_ (0.1M). A 5 ml water sample was collected from all beakers 1- and 7-hours following pH adjustment for the measurement of Na^+^ and Ca^2+^ concentrations. Net ion fluxes were determined by subtracting the ion concentrations at the start of the exposure (1 h sample) from those at the end (7 h sample). Following experimentation, all larvae were lightly blotted dry, weighed and returned to their holding tanks.

**Table 1.** Ion concentrations (in μmol L^-1^) in three [Ca^2+^]_e_e ASW treatments used for acclimation and exposure. Distilled water plus: 5 μmol L^-1^ [Ca^2+^]_e_ = (in μmol L^-1^) 5 CaCl_2_ .7H_2_O, 40 MgSO_4_ .7H_2_O, 490 NaCl, 23 NaOH, 20 KCl; 50 μmol L^-1^ [Ca^2+^]_e_ = (in μmol L^-1^) 50 CaCl_2_ .7H_2_O, 40 MgSO_4_ .7H_2_O, 400 NaCl, 113 NaOH, 20 KCl; and 250 μmol L^-1^ [Ca^2+^]_e_ = (in μmol L^-1^) 250 CaCl_2_ .7H_2_O, 40 MgSO_4_ .7H_2_O, 513 NaOH, 20 KCl.

| $[\text{Ca}^{2+}]$ | $[\text{Na}^+]$ | $[\text{Cl}^-]$ | $[\text{Mg}^{2+}]$ | $[\text{K}^+]$ |
| --- | --- | --- | --- | --- |
| 5 | 513 | 520 | 40 | 20 |
| 50 | 513 | 520 | 40 | 20 |
| 250 | 513 | 520 | 40 | 20 |

### Effects of chronic exposure to high or low Ca^2+^ levels on whole animal Na^+^ and Ca^2+^ fluxes at low pH (experiment 2)

To assess the effect of chronic exposure (acclimation) to high or low [Ca^2+^] on whole animal net Na^+^ and Ca^2+^ fluxes following acute exposure to low pH, *L. terraereginae* larvae (n = 24; Gosner stages 26-38) were transferred to tanks containing either 5 or 250 μmol L^-1^ [Ca^2+^] ASW four weeks prior to sampling. Control larvae were maintained in 50 µmol L^-1^ ASW. Thirty minutes prior to testing, larvae were placed into individual 200 ml glass beakers containing 50 ml of ASW containing the same [Ca^2+^] as their holding tanks (n = 12 per treatment). Water pH in the beakers in each treatment was then acutely lowered to pH 3.5 through the addition of 0.1M H_2_SO_4_. Water samples were then collected from all beakers at 1 h and 7 h and analysed as detailed above for Na^+^ and Ca^2+^ concentrations.

### Effects of Ca^2+^ uptake inhibition on whole animal Na^+^ and Ca^2+^ fluxes acclimated to low pH (experiment 3)

Ruthenium red was used to inhibit apical Ca^2+^ transport since it does not penetrate TJs and is commonly used as a histochemical marker of the barrier formed by epithelial TJs (González-Mariscal et al., 1989; West et al., 2002). *L. terraereginae* larvae (n = 12; Gosner stages 26-38) were randomly allocated to an isolated 5 L tank containing 50 µmol L^-1^ [Ca^2+^]_e_ ASW at pH 3.5 and maintained for four weeks as detailed in experiment 1. Thirty minutes prior to testing, larvae (n = 12) were placed into individual 200 ml glass beakers containing 50 ml of 50 μmol L^-1^ [Ca^2+^]_e_ ASW at pH 3.5. Ruthenium red was added to half of the beakers to a concentration proven to induce an approximately half-maximal response in ECaC activity *in vitro* (10 μmol L^-1^; Hoenderop et al., 2001). Water samples were collected at 1 and 7 h post exposure and analysed as described above. Larvae were then removed from beakers, blotted dry and weighed.

### Gene expression of Ca^2+^ transport and adherens junction proteins

To assess whether [Ca^2+^]_e_ exposure and low pH influences the expression of branchial Ca^2+^ transport proteins and E-cadherin, *L. terraereginae* larvae (n = 36; Gosner stages 26-38) were randomly allocated to six isolated 5 L tanks (n = 6 per tank) containing 5, 50, and 250 μmol L^-1^ [Ca^2+^]_e_ ASW. Water pH was maintained at pH 6.5 (n = 18) or reduced to pH 3.5 (n = 18). Larvae were maintained under these conditions for four weeks. *L. terraereginae* larvae were then euthanised by immersion in 0.25 mg L^-1^ buffered MS222 (Ramlochansingh et al., 2014), and pithing. Both branchial baskets were dissected free and stored in RNAlater (Ambion Inc.) at 4°C for 24 hours, before being moved into a -20°C freezer. Total RNA was extracted from *L. terraereginae* gills using a RNeasy Mini Kit following the manufacturer’s instructions (Qiagen, Valencia, CA). Total RNA was eluted from the silicon spin column in ultrapure water and its concentration quantified using a Qubit fluorometer (ThermoFisher Scientific, Waltham, USA). Any residual genomic DNA contamination was removed, and RNA was reverse transcribed using an iScript gDNA Clear cDNA Synthesis Kit (Bio-Rad, Hercules, CA) following the manufacturers guidelines. Appropriate no reverse transcriptase controls were generated by replacing reverse transcriptase with water.

The transcripts for target genes (ECaC, Calb1, PMCA, NCX, E-Cadherin) and house-keeping genes (β-actin, GAPDH, RPS, TUB) were identified using an in-house *L. terraereginae* transcriptome using homologous sequences from other amphibians as the reference query. Reference sequences were compared against the *L. terraereginae* transcriptome using the ‘blastn’ tool in Galaxy Australia (Jalili et al., 2020). Putative *L. terraereginae* gene sequences were then compared against the National Centre for Biotechnology Information (NCBI) database using the ‘blastn’ tools default parameters to confirm their identity. PrimerQuest (Integrated DNA Technologies, Caraville, IA) was used to design specific qPCR primers (Table S1). All primer pairs were evaluated for specificity and to ensure that they produced only a single band of the appropriate length using MyTaq DNA Polymerase (Bioline, Alexandria, NSW, Australia) and agarose gel electrophoresis.

Quantitative PCR assays were conducted using iTaq™ Universal SYBR® Green Supermix (Bio-rad, Hercules, CA) in a Mini Opticon detection system (MJ Mini Cycler; Bio-rad Laboratories Inc., Hercules, CA). Samples were analysed in triplicate and each plate included appropriate no template controls. No reverse transcriptase controls were assessed independently for each biological sample to confirm the absence of genomic DNA contamination. Cycling parameters were as follows: 95°C for 1 min followed by 40 cycles of 95°C for 15 sec and 60°C for 30 sec. Melt (dissociation) curves (65-90°C) were conducted after each run. Reaction efficiencies were calculated using a serially diluted pooled cDNA standard. All PCR efficiencies were greater than 90% with an R^2^ of over 0.99. All assays produced unique and single peak dissociation curves. Data were exported to MS Excel using Bio-Rad CXF Manager (version 3.1, Bio-Rad). The stability of the candidate house-keeping genes was calculated using the geNorm algorithm via the NormqPCR package (Perkins et al., 2012). All four genes were found to be highly stable across the six treatment groups and met the criteria for designation as appropriate housekeeping genes. A combination of housekeeping genes was used as a pseudo-housekeeper (Rocha-Martins et al., 2012; Vandesompele et al., 2002). The geNorm algorithm revealed the two most stable housekeeping genes (GAPDH and RPS) combined were an effective housekeeping control. To combine the housekeeping genes, the geometric mean of the raw amplification threshold (Ct) values and the reaction efficiencies for GAPDH and RPS were calculated for use in analyses. The following calculations for target gene expression were conducted following Pfaffl (2001). To account for differing reaction efficiencies between primers, adjusted Ct values for the pseudo-housekeeper and the genes of interest were calculated using the following formula:

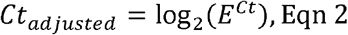

where *E* is the reaction efficiency for the gene and *Ct* is the raw Ct value of the sample. Delta (Δ) Ct was calculated for each sample for statistical analyses:

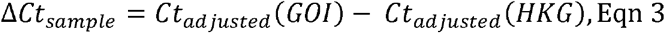

where *Ct*_*adjusted*_*(GOI)* are the adjusted Ct values for the gene of interest, and *Ct*_*adjusted*_*(HKG)* are the adjusted Ct values for the combined housekeeping genes. ΔCt in each target gene was quantified as fold-change relative to the expression of a reference group (the 50 μmol L^-1^ [Ca^2+^]_e_ at pH 6.5 group).

### Statistical analyses

All analyses were conducted in the R statistical environment (R Computing Team, 2021). α was set at 0.05 for all statistical tests. Models were two-tailed and assumed a Gaussian error structure and data satisfied assumptions of hypothesis tests. The effects of body size on rates of ion flux were accounted for by considering wet body mass as a covariate in statistical models.

Differences in net Na^+^ and Ca^2+^ flux between treatment groups were tested by fitting analysis of covariance (ANCOVA) models using the *car* package (Fox and Weisberg, 2018). For experiment 1, a two-way ANCOVA was fitted using exposure [Ca^2+^]_e_ and test pH as independent variables. To determine whether acclimation to low or high Ca^2+^ affected Na^+^ and Ca^2+^ fluxes at low pH in experiment 2, larvae reared at 50 μmol L^-1^ [Ca^2+^]_e_ and exposed to low pH at 5 and 250 μmol L^-1^ [Ca^2+^]_e_ from experiment 1 were compared with larvae reared at 5 or 250 μmol L^-1^ [Ca^2+^]_e_ and exposed to low pH at 5 and 250 μmol L^-1^ [Ca^2+^]_e_. A two-way ANCOVA was fitted modelling test [Ca^2+^]_e_ and rearing [Ca^2+^]_e_ (equimolar to test [Ca^2+^]_e_ versus 50 μmol L^-1^ [Ca^2+^]_e_ control) as independent variables. For experiment 3, a one-way ANCOVA was fitted with RR treatment as the independent variable. For all experiments, one-sample Student’s t-tests were used to test if Na^+^ and Ca^2+^ fluxes in control groups were significantly different from zero suggesting a departure from homeostasis. Net Na^+^ and Ca^2+^ fluxes were assessed in separate models. Post hoc analyses for all ion flux experiments were conducted with the *emmeans* package (Lenth, 2022) using the Tukey method of p value adjustment for multiple comparisons. All data reported are estimated marginal means adjusted for the effect of the covariate.

Differences in target gene expression between treatment groups was analysed by comparing ΔCt values in analysis of variance models using the *car* package (Fox and Weisberg, 2018). A two-way analysis of variance model was fitted using acclimation [Ca^2+^]_e_ and pH as independent variables. This model was fitted for all genes of interest. *Post hoc* analyses were conducted using Tukey’s Honestly Significant Difference test for multiple comparisons.

## RESULTS

### Effect of acute [Ca^2+^]_e_ exposure on Na^+^ and Ca^2+^ fluxes at low pH

Baseline net Na^+^ and Ca^2+^ fluxes in control larvae (pH 6.5, 50 μmol L^-1^ [Ca^2+^]_e_ acclimated) at pH 6.5 were close to zero, although there was a small but significant net loss of Na^+^ (T_5_ = -5.21, p < 0.01; Fig. 2). Net Ca^2+^ fluxes in control larvae were not significantly different from zero. There was a significant interaction between pH and [Ca^2+^]_e_ on net Na^+^ efflux in *L. terraereginae* larvae (F_29_,_2_ = 6.18, *p* < 0.01). Irrespective of [Ca^2+^]_e_, all larvae exposed acutely to pH 3.5 water experienced a substantial net loss of Na^+^ indicating that Na^+^ efflux rates were considerably higher than Na^+^ uptake rates. The magnitude of the effect of acute low pH exposure on Na^+^ flux was greatest in larvae exposed to 5 μmol L^-1^ [Ca^2+^]_e_. There was less effect of low pH on Na^+^ efflux in larvae exposed to 50 μmol L^-1^ (*t*_29_ = 3.2, *p* < 0.01) and 250 μmol L^-1^ [Ca^2+^] (*t*_29_ = 2.56, p = 0.041). There was no effect of pH or [Ca^2+^] on net Ca^2+^ fluxes.

**Fig. 2.**
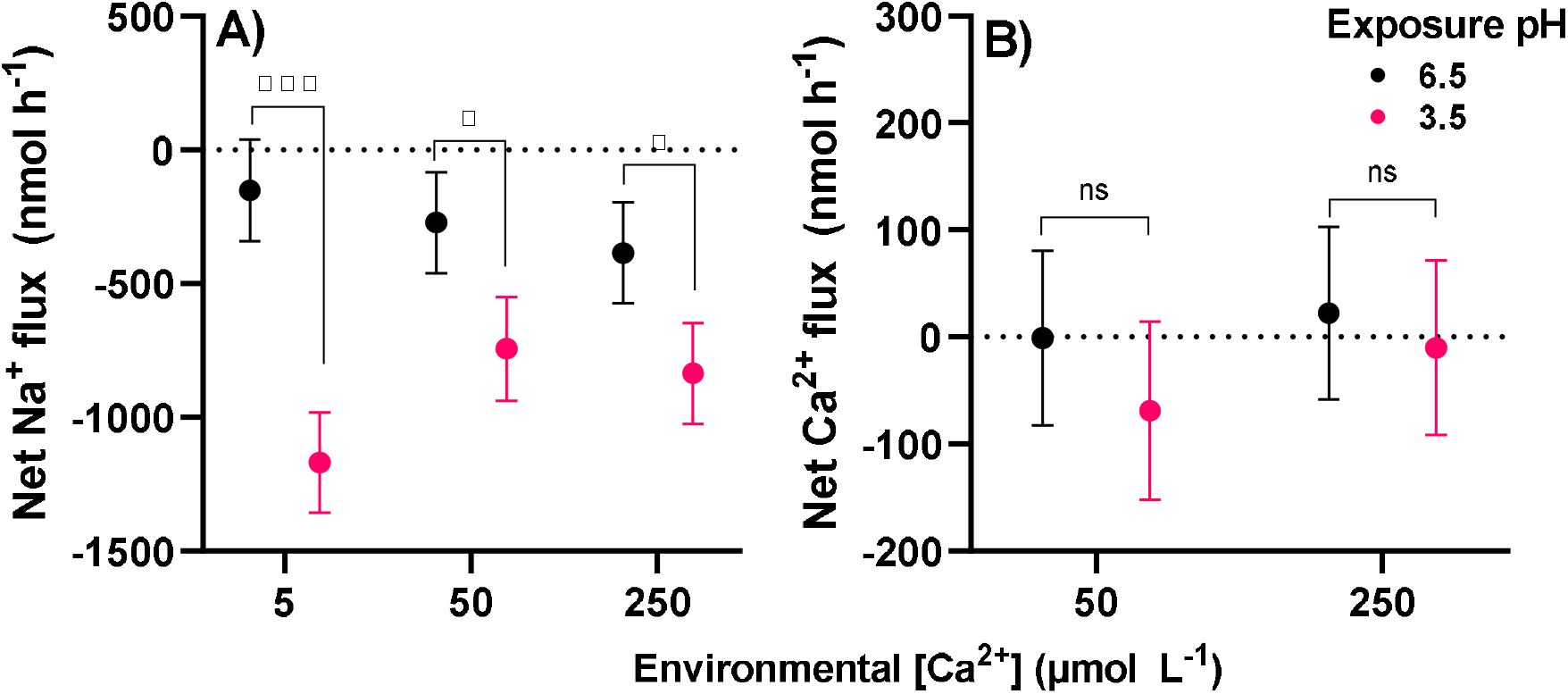
Net Na^+^ (A) and Ca^2+^ (B) fluxes in response to low pH following acute (experiment 1) exposure to varying environmental Ca concentrations ([Ca^2+^]_e_) in *L. terraereginae larvae*. N = 12 larvae within each treatment group in experiment 1 were reared in 50 μmol L^-1^ [Ca^2+^]_e_ and acutely exposed to low pH at either 5, 50, and 250 μmol L^-1^ [Ca^2+^]_e_. Points indicate the estimated marginal means adjusted for body mass, and error bars represent 95% confidence intervals for the fitted models. Positive net flux indicates a net ionic influx, whereas negative net flux indicates a net ionic efflux.

### Effect chronic [Ca^2+^]_e_ exposure on Na^+^ and Ca^2+^ fluxes at low pH

Relative to larvae simultaneously exposed to low pH and acute alterations to [Ca^2+^]_e_ 4-weeks of acclimation to both low and high calcium levels reduced the impact of acute low pH exposure on net Na^+^ fluxes (F, = 9.59, *p* < 0.01; Fig. 3). Similarly, there was a significant effect of acclimation to 250 μmol L^-1^ [Ca^2+^]_e_ on net Ca^2+^ fluxes, with larvae acclimated to high [Ca^2+^]_e_ experiencing a significant net Ca^2+^ influx (compared with larvae in 50 μmol L^-1^ [Ca^2+^]_e_) (F_19,1_, = 12.56, *p* < 0.01).

**Fig. 3.**
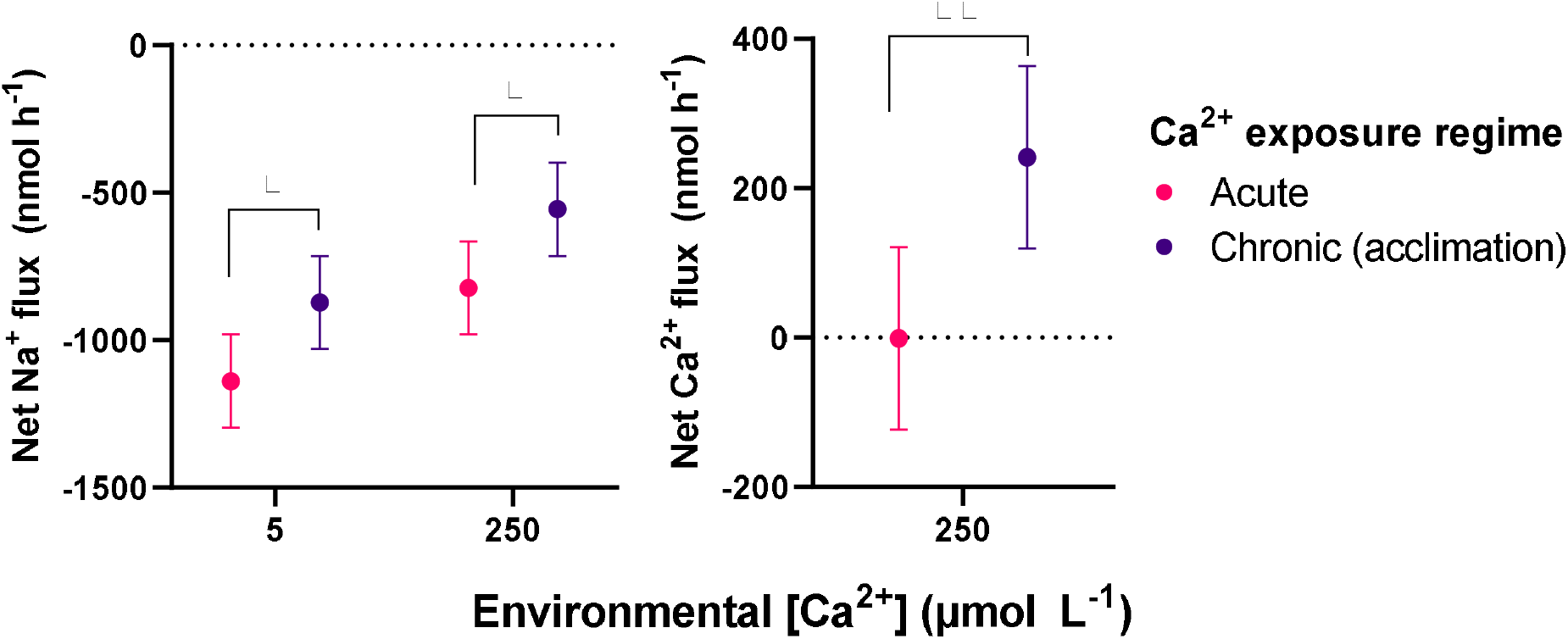
The effect of acute or chronic exposure to low and high environmental Ca^2+^ concentrations ([Ca^2+^]_e_) on whole animal Na^+^ and Ca^2+^ fluxes following acute exposure to low pH. N = 12 larvae within each treatment group in experiment 1 reared in 50 μmol L^-1^ [Ca^2+^]_e_ and acutely exposed to low pH at either 5 or 250 μmol L^-1^ [Ca^2+^]_e_ were compared to n = 12 larvae within each treatment group in experiment 2 reared in 5 or 250 μmol L^-1^ [Ca^2+^]_e_ and acutely exposed to low pH at equimolar [Ca^2+^]. Points indicate the estimated marginal means adjusted for body mass, and error bars represent 95% confidence intervals for the fitted models. Positive net flux indicates a net ionic influx, whereas negative net flux indicates a net ionic efflux.

### Effect of the inhibition of apical Ca^2+^ uptake on Na^+^ and Ca^2+^ fluxes in larvae acclimated to low pH

In *L. terraereginae* larvae reared from hatching at pH 3.5 and with 50 μmol L^-1^ [Ca^2+^]_e_ ASW, baseline net Na^+^ efflux was small but significantly lower than zero (T_5_ = -4.1833, p < 0.01; Fig. 4). Larvae had a baseline net Ca^2+^ influx that was slightly but significantly higher than zero (*T*_5_ = 2.7916, *p* = 0.038). Acute exposure of *L. terraereginae* larvae to ruthenium red resulted in a substantial increase in both net Na^+^ efflux (F_9,1_ = 71.174, *p* = <0.001) and net Ca^2+^ efflux (F_9,1_ = 38.14, *p* < 0.001) over the 6 h exposure period.

**Fig. 4.**
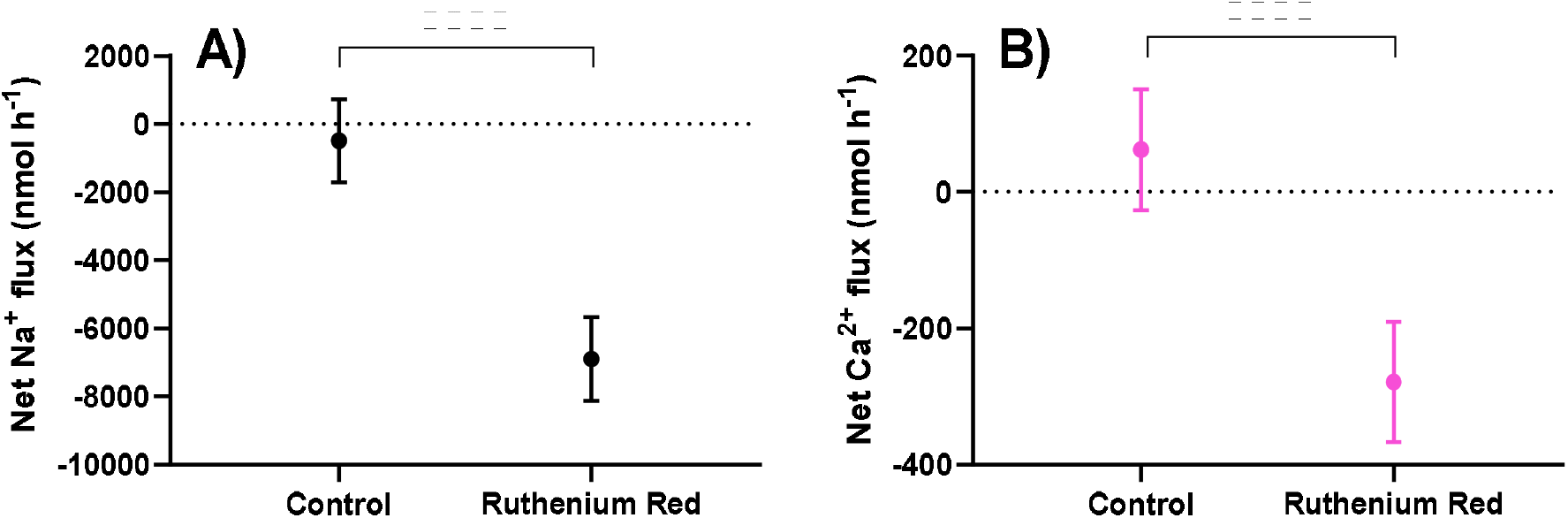
Net Na^+^ (A) and Ca^2+^ (B) fluxes in N = 12 *L. terraereginae* larvae within each group reared at pH 3.5 with 50 μmol L^-1^ environmental Ca^2+^ concentration ([Ca^2+^] _e_), acutely exposed 10 μmol L^-1^ RR at pH 3.5. Points indicate the estimated marginal means adjusted for mass, and error bars represent 95% confidence intervals for the fitted model. Net Na^+^ and Ca^2+^ fluxes were significantly higher in *L. terraereginae* acutely exposed to 10 μmol L^-1^ ruthenium red for 6 h.

### Gene expression

The expression of key Ca^2+^ transport proteins and E-cadherin in the gills of *L. terraereginae* was compared across larvae reared at pH both 6.5 and pH 3.5 in low, moderate or high [Ca^2+^]_e_. There was no significant effect of acclimation pH or [Ca^2+^]_e_ on the gene expression of branchial ECaC, PMCA, or NCX channels (Fig. 5). However, branchial calbindin expression was significantly higher in larvae reared at pH 3.5 than in pH 6.5-reared larvae (F_1,24_ = 5.640, p = 0.026) but was not influenced by [Ca^2+^]_e_. Branchial E-cadherin mRNA expression was significantly higher in larvae reared under high [Ca^2+^]_e_ levels (F_2,24_ = 5.177, *p* = 0.0135) but was unaffected by acclimation pH.

**Fig. 5.**
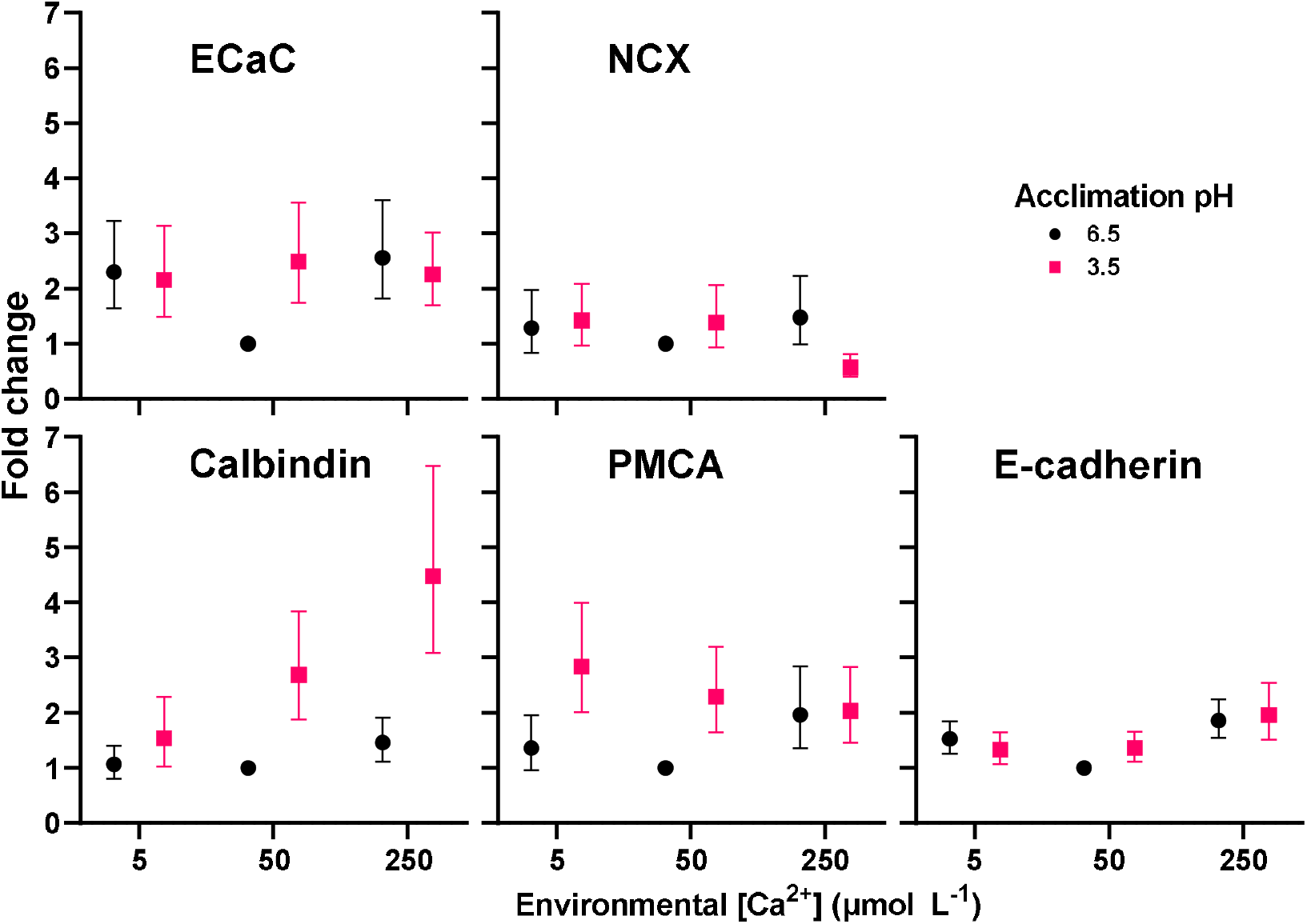
Expression of five key Ca^2+^ transport genes in N = 6 *L. terraereginae* larvae per group reared in circumneutral and low pH ASW containing 5, 50, and 250 μmol L^-1^ environmental Ca^2+^ concentration ([Ca^2+^]_e_). Change in expression is represented as fold change made relative to the pH 6.5, 50 μmol L^-1^ [Ca^2+^]_e_ control treatment group. Points indicate mean fold change and error bars indicate standard error using a Taylor series (Weng et al., 2006).

## DISCUSSION

Highly acidic waters pose a major threat to transcellular Ca^2+^ uptake pathways, which may play a role in the acute and potentially fatal branchial Na^+^ loss experienced by acid-sensitive aquatic animals exposed to low pH. Conversely, acid tolerant animals may employ a suite of mechanisms that enable them to protect Ca^2+^ uptake capacity which in turn allows them to resist the negative effects of low pH on junctional integrity. Consistent with our hypothesis, acute exposure to low pH water in the presence of low [Ca^2+^]_e_ increased net Na^+^ efflux, but not net Ca^2+^ fluxes in acid-tolerance *L. terraereginae* larvae. Provision of additional [Ca^2+^]_e_ reduced net Na^+^ efflux rates, but this effect was saturable. Acclimation to both low and high [Ca^2+^]_e_ improved the resistance of larvae to Na^+^ efflux at low pH and resulted in an increased net Ca^2+^ influx. Inhibition of apical epithelial calcium uptake by ruthenium red resulted in the complete loss of tolerance to low pH in larvae acclimated to low pH water consistent with our hypothesis that acclimation to low pH involves the protection of Ca^2+^ uptake capacity. Acclimation to low pH water increased branchial gene expression of the intracellular Ca^2+^ transport protein calbindin, consistent with a role for increased transcellular Ca^2+^ trafficking in the tolerance of low pH water. These results establish a significant role for the security of Ca^2+^ uptake capacity in the tolerance of *L. terraereginae* larvae living in highly acidic waters.

Acid-naive *L. terraereginae* larvae reared at circumneutral pH and acutely exposed to low pH had a greater rate of net Na^+^ efflux than larvae reared and tested at pH 6.5, consistent with an acute negative effect of low pH on intercellular junction integrity. The high rate of net Na^+^ efflux was substantially reduced in animals acclimated to low pH consistent with an acid-tolerant phenotype. In acid-naïve L. terraereginae, the high rate of net Na^+^ efflux was exacerbated in the presence of low [Ca^2+^]_e_, indicating a protective effect of [Ca^2+^]_e_ on epithelial junction integrity. This is consistent with previous studies showing that [Ca^2+^]_e_ plays a significant role in determining Na^+^ efflux rates with low pH exposure in a range of fish and amphibian larvae (Cummins, 1988; Freda and Dunson, 1984; Freda et al., 1991; Gascon et al., 1987; Gonzalez and Dunson, 1989; Gonzalez et al., 1998; Kullberg et al., 1993; Kumai et al., 2011; McDonald and Rogano, 1986; McDonald et al., 1983; Meyer et al., 2010; Riesch et al., 2015). However, environmental Ca^2+^ was only beneficial for controlling Na^+^ efflux up to a point: exposure of larvae to 250 μmol L^-1^ [Ca^2+^]_e_ did not further reduce net Na^+^ fluxes beyond that of larvae exposed to control (50 μmol L^-1^) levels. This may indicate the influence of elevated Ca^2+^ on Na^+^ efflux is saturable, and 50 μmol L^-1^ [Ca^2+^]_e_ completely saturates the gill epithelium. This is consistent with Meyer (2004) who demonstrated that 80 μmol L^-1^ [Ca^2+^]_e_ reduced Na^+^ efflux during low pH exposure in *L. terraereginae* larvae, however raising [Ca^2+^]_e_ to 400 μmol L^-1^ had no further effects.

The protective effect of [Ca^2+^]_e_ on branchial junction permeability has been attributed to extracellular Ca^2+^ bound to the gill epithelium, specifically to the TJ (Freda and McDonald, 1988; Gonzalez and Dunson, 1989; Mcdonald, 1983; McWilliams, 1983; Reid et al., 1991; Yu et al., 2010). However, these studies did not account for the possibility that low pH affects junctional stability by impairing transcellular Ca^2+^ uptake pathways. Exposure of *L. terraereginae* larvae to water containing 10 μmol L^-1^ of ruthenium red, a potent ECaC channel inhibitor (Nilius et al., 2001), resulted in a large (7500 – 17500 nmol h^-1^) increase in net Na^+^ efflux. This suggests that the inhibition of the transcellular Ca^2+^ uptake pathway has an immediate and severe effect on junctional stability leading to increased junctional permeability to Na^+^. Similar findings have been observed in studies of goldfish and tetras exposed to La^3+^, a Ca^2+^ analogue that, unlike ruthenium red, also has the capacity to penetrate TJs and interact with AJs directly (Eddy and Bath, 1979; Gonzalez et al., 1997; Lacaz-Vieira and Marques, 2004). The inhibition of transcellular Ca^2+^ uptake using ruthenium red has not been performed previously on whole animals but does show that inhibition of Ca^2+^ uptake can have catastrophic impacts on homeostasis consistent with the loss of junctional stability. We posit that the inhibition of apical Ca^2+^ uptake disrupts AJ stability with consequences for the maintenance of TJ stability. Since ECaC activity is inhibited by low pH (Vennekens et al., 2001), we propose that a loss of Ca^2+^ uptake capacity following acute exposure to low pH may underpin the loss of junctional stability and resulting Na+ efflux in acid sensitive organisms. Conversely, adaptations which counter ECaC inhibition at low pH may protect Ca^2+^ uptake capacity in acidophilic species. While the effects of ruthenium red on Na^+^ efflux in acid-acclimated larvae were rapid and extreme, an investigation of its effects on junctional morphology would be needed to demonstrate that high Na^+^ losses were the result of junctional disruption and not the inhibition of other Na^+^ and Ca^2+^ transport pathways. While ruthenium red is a potent inhibitor of ECaC, it can also affect other Ca^2+^ transport proteins which may affect the maintenance of ion balance (Hajnóczky et al., 2006; Vincent and Duncton, 2011).

Unlike Na^+^ fluxes, acute exposure to low pH water did not affect net Ca^2+^ fluxes when larvae were acutely exposed to high or control [Ca^2+^]_e_. In both circumneutral and low pH water, Ca^2+^ fluxes were not significantly different from zero suggesting that rates of influx and efflux were balanced. This was unexpected as low pH has been shown to inhibit Ca^2+^ uptake (ECaC activity) in multiple cell lines (Bindels et al., 1994; Hoenderop et al., 1999; Vennekens et al., 2001). If low pH also inhibited ECaC activity in L. terraereginae, it should have manifested as a net increase in the rate of Ca^2+^ loss. In fact, *L. terraereginae* larvae reared in high [Ca^2+^]_e_ and acutely exposed to low pH experienced a net Ca^2+^ influx (approx. 250 nmol h^-1^) compared to animals reared at control [Ca^2+^]_e_ levels. In low pH reared larvae, there was an apparent net increase in Ca^2+^ uptake. These influxes strongly suggests that ECaC in the branchial epithelium of *L. terraereginae* larvae is not substantially inhibited by protonation, and that transcellular environmental Ca^2+^ uptake is maintained at low pH and may play a role in facilitating acid tolerance in L. terraereginae. This may highlight an adaptation for the prevention of Ca^2+^ uptake inhibition at low pH and could be linked to the expression of a less pH sensitive ECaC isoform. Consistent with this hypothesis, inhibition of apical Ca^2+^ uptake by ruthenium red did result in a large increase in net Ca^2+^ efflux in larvae. Clearly, the disturbance of transcellular Ca^2+^ uptake has serious implications for the maintenance of transepithelial resistance via the loss of junction stability at low pH.

The maintenance of transcellular Ca^2+^ transport is potentially important in promoting acid tolerance of acidophilic animals. Ca^2+^ might influence tolerance of low pH via association with the Ca^2+^-dependent AJ, which is directly responsible for the stability of the TJ and thus epithelial permeability, a major factor in preventing Na^+^ at low pH in acidophilic species (Gumbiner et al., 1988; Kwong et al., 2014; Watabe-Uchida et al., 1998). Lowering intracellular [Ca^2+^] in MDCK cells has been shown to interfere with the formation of TJs (Stuart et al., 1996). Ca^2+^ also employs many signalling functions such as hormone regulation (Clapham, 2007; D’Souza-Li, 2006), which may potentially alter the expression of genes involved in maintaining junctional integrity at low pH. Calbindin and other intracellular Ca^2+^-binding proteins function to buffer intracellular Ca^2+^ concentrations by facilitating the basolateral extrusion of Ca^2+^ taken up across the apical membrane (Christakos et al., 1989). The finding that calbindin mRNA was upregulated in pH 3.5 reared *L. terraereginae* larvae is suggestive of increased transcellular Ca^2+^ movements in the gill epithelium of *L. terraereginae* larvae acclimated to low pH. Interestingly, cytosolic Ca^2+^ is critical in regulating ECaC activity (Hoenderop et al., 1999). This finding is consistent with the idea that transcellular Ca^2+^ transport is involved in acid tolerance to some degree and that Ca^2+^ uptake is not inhibited by low pH in *L. terraereginae* larvae.

In contrast to our hypotheses, mRNA expression of key transcellular Ca^2+^ transport proteins ECaC, NCX and PMCA were not influenced by environmental pH or [Ca^2+^]_e_ in *L. terraereginae* larvae despite the observation that acclimation to low pH was accompanied by an apparent net increase in Ca^2+^ uptake. If this effect was indeed the result of an increase in Ca^2+^ uptake and not a reduction in efflux, then it is possible that it was facilitated by an increase in the activity of the existing channels as opposed to the de novo production of more channels. Likewise, the lack of increase in E-cadherin mRNA expression with chronic low pH exposure suggests that E-cadherin function was unaffected by low pH. E-cadherins bind Ca^2+^ from the extracellular environment; our data suggest that maintenance of the transcellular Ca^2+^ transport pathway allows for the maintenance of favourable Ca^2+^ concentrations in the extracellular space to prevent E-cadherin disfunction. Consistent with this idea, increased expression of calbindin mRNA in acid-acclimated larvae provides some evidence that increased transcellular Ca^2+^ transport plays a role in promoting extreme acid tolerance. Increased Ca^2+^ uptake at the apical membrane may be evidenced by increased intracellular Ca^2+^ shuttling rates (and an associated increased abundance of calbindin proteins) to maintain Ca^2+^-dependent junction dynamics in low pH environments. The finding that low pH acclimated larvae upregulated calbindin mRNA and had a net Ca^2+^ influx supports a role for increased transcellular [Ca^2+^]_e_ movement in facilitating acid tolerance in *L. terraereginae* larvae. However, mRNA expression levels do not always correlate well with actual levels of protein expression, so care must be taken when interpreting mRNA expression patterns in the absence of corresponding protein expression levels. Differential post-translational processing of mRNA and other factors can be responsible for the low correlation between an organism’s transcriptome and its proteome (Ghazalpour et al., 2011; Marguerat et al., 2012). Although we cannot rule out a paracellular route for the uptake of Ca^2+^ into the extracellular space, studies in fish suggests that more than 97% of branchial (gills) Ca^2+^ uptake is active (transcellular) (Flik et al., 1995). The hormonal control of Ca^2+^ transport protein function and abundance is a potentially overlooked factor in understanding how branchial Ca^2+^ transport is influenced by extracellular pH and [Ca^2+^]_e_ in L. terraereginae.

This study showed that environmental Ca^2+^ has a protective effect on the control of Na^+^ efflux at low pH in *L. terraereginae* larvae. Furthermore, it demonstrated that larvae have a capacity for acclimation to low pH via changes in Na^+^ and Ca^2+^ fluxes, which appear to involve the transcellular pathway (i.e. increased calbindin mRNA upregulation). Given that *L. terraereginae* larvae can maintain or increase Ca^2+^ uptake at low pH suggests that protonation of the branchial epithelium likely does not outwardly inhibit Ca^2+^ uptake. However, we only examined net Na^+^ and Ca^2+^ fluxes which do not reveal details about the discrete behaviour of uptake and efflux pathways. The use of radioactive isotopes or fluorescent ion analogues could be used to better resolve changes in influx and efflux pathways and how they contribute to net ion fluxes. Inhibition of apical Ca^2+^ uptake by ruthenium red strongly supports a role for the maintenance of transcellular Ca^2+^ uptake in the control of branchial junction stability at low pH in *L. terraereginae* but does not reveal the specific site of action for Ca^2+^ in preventing Na^+^ loss at low pH. Using intracellular Ca^2+^ markers to track Ca^2+^ movement during acclimation to low pH may help to elucidate this mechanism. As *L. terraereginae* larvae are exceptionally acid tolerant, their ability to maintain Ca^2+^ uptake in very soft and acidic waters may be a unique adaptation. Comparing transcellular Ca^2+^ transport capabilities with other acid-sensitive species might reveal the mechanistic adaptations employed by *L. terraereginae* in the maintenance of epithelial stability at low pH. The current study highlights a role for transcellular Ca^2+^ transport and the prevention of Ca^2+^ uptake inhibition by low pH in the extreme acid tolerance of *L. terraereginae* larvae.

## ACKNOWLEDGEMENTS

Funding for this research was provided by the Australian Research Council (DP150101571) to CEF and was done under the UQ Animal Ethics Committee (Approval number: SBS/460/14/ARC). The authors thank Dr Edward Meyer for useful discussions during the project, Dr Nicholas Wu and Callum McKercher for assistance with gene expression studies.

## DATA AVAILABILITY

The complete datasets and R scripts used for analysing the data is publicly available at UQ eSpace (https://doi.org/10.14264/18c7301)

## AUTHOR CONTRIBUTIONS

Conceptualisation: RLC, CH, CEF; Methodology: CH, RLC, CEF; Validation: CH; Formal analysis: CH; Investigation: CH; Resources: CEF; Data curation: CH; Writing – original draft: CH; Writing – review & editing: CH, RLC, CEF; Visualisation: CH; Supervision: RLC, CEF; Project administration: RLC, CEF; Funding acquisition: RLC, CEF.

## REFERENCES

Baldwin, G. F. and Bentley, P. J. (1980). Calcium metabolism in bullfrog tadpoles (Rana catesbeiana). J. Exp. Biol. 88, 357–365.

Bayly, I. A. (1964). Chemical and biological studies on some acidic lakes of east Australian sandy coastal lowlands. Mar. Freshw. Res. 15, 56–72.

Bhat, M., Toledo-Velasquez, D., Wang, L. Y., Malanga, C. J., Ma, J. K. H. and Rojanasakul, Y. (1993). Regulation of tight junction permeability by calcium mediators and cell cytoskeleton in rabbit tracheal epithelium. Pharm. Res. An Off. J. Am. Assoc. Pharm. Sci. 10, 991–997.

Bindels, R. J. M., Hartog, A., Abrahamse, S. L. and Van Os, C. H. (1994). Effects of pH on apical calcium entry and active calcium transport in rabbit cortical collecting system. Am. J. Physiol. - Ren. Fluid Electrolyte Physiol. 266, F620–F627.

Brown, R. C. and Davis, T. P. (2002). Calcium modulation of adherens and tight junction function: A potential mechanism for blood-brain barrier disruption after stroke. Stroke 33, 1706–1711.

Cai, Q., Chandler, J. S., Wasserman, R. H., Kumar, R. and Penniston, J. T. (1993). Vitamin D and adaptation to dietary calcium and phosphate deficiencies increase intestinal plasma membrane calcium pump gene expression. Proc. Natl. Acad. Sci. U. S. A. 90, 1345–1349.

Chasiotis, H., Kolosov, D., Bui, P. and Kelly, S. P. (2012). Tight junctions, tight junction proteins and paracellular permeability across the gill epithelium of fishes: A review. Respir. Physiol. Neurobiol. 184, 269–281.

Christakos, S., Gabrielides, C. and Rhoten, W. B. (1989). Vitamin D-dependent calcium binding proteins: Chemistry, distribution, functional considerations, and molecular biology. Endocr. Rev. 10, 3–26.

Clapham, D. (2007). Calcium signaling. Cell 131, 1047–1058.

Cummins, C. P. (1988). Effect of calcium on survival times of Rana temporaria embryos at low pH. Funct. Ecol. 2, 397–302.

D’Souza-Li, L. (2006). The calcium-sensing receptor and related diseases. Arq. Bras. Endocrinol. Metabol. 50, 628–639.

Dalziel, T. R. K., Morris, R. and Brown, D. J. A. (1986). The effects of low pH, low calcium concentrations and elevated aluminium concentrations on sodium fluxes in brown trout, Salmo Trutta. Water Air Soil Pollut. 30, 569–577.

Daye, P. G. and Garside, E. T. (1976). Histopathologic changes in surficial tissues of brook trout, Salvelinus fontinalis (Mitchill), exposed to acute and chronic levels of pH. Can. J. Zool. 54, 2140–2155.

Eddy, F. B. and Bath, R. N. (1979). Effects of lanthanum on sodium and chloride fluxes in the goldfish Carassius auratus. J. Comp. Physiol. 129, 145–149.

Edwards, S. L. and Marshall, W. S. (2012). Principles and Patterns of Osmoregulation and Euryhalinity in Fishes. Fish Physiol. 32, 1–44.

Evans, D. H., Piermarini, P. M. and Potts, W. T. W. (1999). Ionic transport in the fish gill epithelium. J. Exp. Zool. 283, 641–652.

Flik, G. and Verbost, P. M. (1995). Cellular mechanisms in calcium transport and homeostasis in fish. Biochem. Mol. Biol. Fishes 5, 251–263.

Flik, G., Verbost, P. M. and Wendelaar Bonga, S. E. (1995). Calcium transport processes in fishes. In Fish Physiology 141Z: Cellular and Molecular Approaches to Fish Ionic Regulation (ed. Wood, C. M.and Shuttleworth, T. J.), pp. 317–342. San Diego: Academic Press.

Fox, J. and Weisberg, S. (2018). An R Companion to applied regression. 3rd ed. SAGE Publications.

Freda, J. and Dunson, W. A. (1984). Sodium balance of amphibian larvae exposed to low environmental pH. Physiol. Zool. 57, 435–443.

Freda, J. and McDonald, D. G. (1988). Physiological correlates of interspecific variation in acid tolerance in fish. J. Exp. Biol. 136, 243–258.

Freda, J., Sanchez, D. A. and Bergman, H. L. (1991). Shortening of branchial tight junction acid-exposed rainbow trout (Oncorhynchus mykiss). Can. J. Fish. Aquat. Sci. 48, 2028–2033.

Gascon, C., Planas, D. and Moreau, G. (1987). The interaction of pH, calcium, and aluminium concentrations on the survival and development of wood frog (Rana sylvatica) eggs and tadpoles. Ann. La Soc. R. Zool. Belgique 117, 189–199.

Ghazalpour, A., Bennett, B., Petyuk, V. A., Orozco, L., Hagopian, R., Mungrue, I. N., Farber, C. R., Sinsheimer, J., Kang, H. M., Furlotte, N., et al. (2011). Comparative analysis of proteome and transcriptome variation in mouse. PLoS Genet. 7, e1001393.

Gonzalez-Mariscal, L., Contreras, R. G., Bolivar, J. J., Ponce, A., Chavez De Ramirez, B. and Cereijido, M. (1990). Role of calcium in tight junction formation between epithelial cells. Am. J. Physiol. 259, C978–C986.

González-Mariscal, L., de Ramirez, B. C., Lázaro, A. and Cereijido, M. (1989). Establishment of tight junctions between cells from different animal species and different sealing capacities. J. Membr. Biol. 107, 43–56.

Gonzalez, R. J. and Dunson, W. A. (1989). Acclimation of sodium regulation to low pH and the role of calcium in the acid-tolerant sunfish Enneacanthus obesus. Physiol. Zool. 62, 977–992.

Gonzalez, R. J., Dalton, V. M. and Patrick, M. L. (1997). Ion regulation in ion-poor acidic water by the blackskirt tetra (Gymnocorymbus ternetzi), a fish native to the Amazon River. Physiol. Zool. 70, 428–435.

Gonzalez, R. J., Wood, C. M., Wilson, R. W., Patrick, M. L., Bergman, H. L., Narahara, A. and Val, A. L. (1998). Effects of water pH and calcium concentration on ion balance in fish of the Rio Negro, Amazon. Physiol. Zool. 71, 15–22.

Gosner, K. (1960). A simplified table for staging anuran embryos and larvae with notes on identification. Herpetologica 16, 183–190.

Gumbiner, B., Stevenson, B. and Grimaldi, A. (1988). The role of the cell adhesion molecule uvomorulin in the formation and maintenance of the epithelial junctional complex. J. Cell Biol. 107, 1575–1587.

Hajnóczky, G., Csordás, G., Das, S., Garcia-Perez, C., Saotome, M., Sinha Roy, S. and Yi, M. (2006). Mitochondrial calcium signalling and cell death: Approaches for assessing the role of mitochondrial Ca^2+^ uptake in apoptosis. Cell Calcium 40, 553–560.

Hartsock, A. and Nelson, W. J. (2008). Adherens and tight junctions: Structure, function and connections to the actin cytoskeleton. Biochim. Biophys. Acta 1778, 660–669.

Hines, H. and Meyer, E. (2011). The frog fauna of Bribie Island: An annotated list and comparison with other Queensland dune islands. Proc. R. Soc. Queensl. 117, 261–274.

Hoenderop, J. G. J., Van Der Kemp, A. W. C. M., Hartog, A., Van Os, C. H., Willems, P. H. G. M. and Bindels, R. J. M. (1999). The epithelial calcium channel, ECaC, is activated by hyperpolarization and regulated by cytosolic calcium. Biochem. Biophys. Res. Commun. 261, 488–492.

Jalili, V., Afgan, E., Gu, Q., Clements, D., Blankenberg, D., Goecks, J., Taylor, J. and Nekrutenko, A. (2020). The Galaxy platform for accessible, reproducible and collaborative biomedical analyses: 2020 update. Nucleic Acids Res. 48, W395–W402.

Kelly, S. P. and Wood, C. M. (2008). Cortisol stimulates calcium transport across cultured gill epithelia from freshwater rainbow trout. Vitr. Cell. Dev. Biol. 44, 96–104.

Khanal, R. C. and Nemere, I. (2008). Regulation of intestinal calcium transport. Annu. Rev. Nutr. 28, 179–196.

Kullberg, A., Bishop, K. H., Hargeby, A., Jansson, M. and Petersen, R. C. (1993). The ecological significance of dissolved organic carbon in acidified waters. Ambio 22, 331–337.

Kumai, Y., Bahubeshi, A., Steele, S. and Perry, S. F. (2011). Strategies for maintaining Na^+^ balance in zebrafish (Danio rerio) during prolonged exposure to acidic water. Comp. Biochem. Physiol. 160, 52–62.

Kwong, R. W. M., Kumai, Y. and Perry, S. F. (2014). The physiology of fish at low pH: The zebrafish as a model system. J. Exp. Biol. 217, 651–662.

Lacaz-Vieira, F. and Marques, M. M. (2004). Lanthanum effect on the dynamics of tight junction opening and closing. J. Membr. Biol. 202, 39–49.

Lenth, R. (2022). Emmeans: Estimated marginal means, aka least-square means. R package version 1.7.3. https://CRAN.R-project.org/package=emmeans.

Ma, T. Y., Tran, D., Hoa, N., Nguyen, D., Merryfield, M. and Tarnawski, A. (2000). Mechanism of extracellular calcium regulation of intestinal epithelial tight junction permeability: Role of cytoskeletal involvement. Microsc. Res. Tech. 51, 156–168.

Marguerat, S., Schmidt, A., Codlin, S., Chen, W., Aebersold, R. and Bähler, J. (2012). Quantitative analysis of fission yeast transcriptomes and proteomes in proliferating and quiescent cells. Cell 151, 671–683.

Matsuo, A. Y. and Val, A. L. (2002). Low pH and calcium effects on net Na^+^ and K^+^ fluxes in two catfish species from the Amazon River (Corydoras: Callichthyidae). Brazilian J. Med. Biol. Res. 35, 361–367.

Mcdonald, D. G. (1983). The interaction of environmental calcium and low pH on the physiology of the rainbow trout, Salmo gairdneri: I. Branchial and renal net ion and H^+^ Fluxes. J. Exp. Biol. 102, 123–140.

McDonald, D. G. and Rogano, M. S. (1986). Ion regulation by the rainbow trout, Salmo gairdneri, in ion-poor water. Physiol. Zool. 59, 318–331.

McDonald, D. G., Walker, R. L. and Wilkes, P. R. H. (1983). The interaction of environmental calcium and low pH on the physiology of the rainbow trout, Salmo Gairdneri II. Branchial ionoregulatory mechanisms. J. Exp. Biol. 102, 141–155.

McDonald, D. G., Ozog, J. L. and Simons, B. P. (1984). The influence of low pH environments on ion regulation in the larval stages of the anuran amphibian, Rana clamitans. Can. J. Zool. 62, 2171–2177.

McWilliams, P. G. (1983). An investigation of the loss of bound calcium from the gills of the brown trout, Salmo trutta, in acid media. Comp. Biochem. Physiol. 74, 107–116.

Meyer, E. (2004). Acid adaptation and mechanisms for softwater acid tolerance in larvae of anuran species native to the “Wallum” of east Australia. PhD thesis, University of Queensland, St Lucia, Australia.

Meyer, E. A., Cramp, R. L. and Franklin, C. E. (2010). Damage to the gills and integument of Litoria fallax larvae (Amphibia: Anura) associated with ionoregulatory disturbance at low pH. Comp. Biochem. Physiol. 155, 164–171.

Meyer, E. A., Franklin, C. E. and Cramp, R. L. (2020). Physiological and morphological correlates of extreme acid tolerance in larvae of the acidophilic amphibian Litoria cooloolensis. J. Comp. Physiol. B 191, 159–171.

Nilius, B., Prenen, J., Vennekens, R., Hoenderop, J. G. J., Bindels, R. J. M. and Droogmans, G. (2001). Pharmacological modulation of monovalent cation currents through the epithelial Ca^2+^ channel ECac1. Br. J. Pharmacol. 134, 453–462.

O’Keefe, E. J., Briggaman, R. A. and Herman, B. (1987). Calcium-induced assembly of adherens junctions in keratinocytes. J. Cell Biol. 105, 807–817.

Perkins, J. R., Dawes, J. M., McMahon, S. B., Bennett, D. L. H., Orengo, C. and Kohl, M. (2012). ReadqPCR and NormqPCR: R packages for the reading, quality checking and normalisation of RT-qPCR quantification cycle (Cq) data. BMC Genomics 13, 296.

Pfaffl, M. W. (2001). A new mathematical model for relative quantification in real-time RT-PCR. Nucleic Acids Res. 29, e45.

Pokutta, S., Herrenknecht, K., Kemler, R. and Engel, J. (1994). Conformational changes of the recombinant extracellular domain of E-cadherin upon calcium binding. Eur. J. Biochem. 223, 1019–1026.

R Computing Team (2021). A language and environment for statistical computing. R Foundation for Statistical Computing, Vienna, Austria. URL https://www.R-project.org/.

Ramlochansingh, C., Branoner, F., Chagnaud, B. P. and Straka, H. (2014). Efficacy of tricaine methanesulfonate (MS-222) as an anesthetic agent for blocking sensory-motor responses in Xenopus laevis tadpoles. PLoS One 9, e101606.

Reid, S. D., McDonald, D. G. and Rhem, R. R. (1991). Acclimation to sublethal aluminum: modifications of metal – gill surface interactions of juvenile rainbow trout (Oncorhynchus mykiss). Can. J. Fish. Aquat. Sci. 48, 1996–2005.

Riesch, R., Tobler, M. and Plath, M. (2015). Extremophile fishes: Ecology, Evolution, and Physiology of Teleosts in Extreme Environments. Switzerland: Springer.

Robinson, G. D. (1993). Effects of Reduced Ambient pH on Sodium Balance in the Red-Spotted Newt, Notophthalmus viridescens. Physiol. Zool. 66, 602–618.

Rocha-Martins, M., Njaine, B. and Silveira, M. S. (2012). Avoiding pitfalls of internal controls: Validation of reference genes for analysis by qRT-PCR and western blot throughout rat retinal development. PLoS One 7, e43028.

Rosseland, B. O. and Staurnes, M. (1994). Physiological mechanisms for toxic effects and resistance to acid water: an ecophysiological and ecotoxicological approach. In Acidification of Freshwater Ecosystems: Implications for the Future (ed. Steinberg, W. C. E.and Wright, R. F.), pp. 227–246. London, UK: John Wiley and Sons.

Schneeberger, E. E. and Lynch, R. D. (1992). Structure, function, and regulation of cellular tight junctions. Am. J. Physiol. 262, L647–661.

Schneeberger, E. E. and Lynch, R. D. (2004). The tight junction: A multifunctional complex. Am. J. Physiol. 286, C1213–C1228.

Shahsavarani, A. and Perry, S. F. (2006). Hormonal and environmental regulation of epithelial calcium channel in gill of rainbow trout (Oncorhynchus mykiss). Am. J. Physiol 291, R1490–R1498.

Stuart, R. O., Sun, A., Bush, K. T. and Nigam, S. K. (1996). Dependence of epithelial intercellular junction biogenesis on thapsigargin-sensitive intracellular calcium stores. J. Biol. Chem. 271, 13636–13641.

Thiers, R. E. and Hviid, K. (1962). Interference-free flame photometry of calcium in serum and urine. Clin. Chem. 8, 35–46.

Vandesompele, J., De Preter, K., Pattyn, F., Poppe, B., Van Roy, N., De Paepe, A. and Speleman, F. (2002). Accurate normalization of real-time quantitative RT-PCR data by geometric averaging of multiple internal control genes. Genome Biol. 3, research0034.1.

Vennekens, R., Prenen, J., Hoenderop, J. G. J., Bindels, R. J. M., Droogmans, G. and Nilius, B. (2001). Modulation of the epithelial Ca^2+^ channel ECaC by extracellular pH. Pflugers Arch. Eur. J. Physiol. 442, 237–242.

Verbost, P. M., Flik, G., Fenwick, J. C., Greco, A. M., Pang, P. K. T. and Wendelaar Bonga, S. E. (1993). Branchial calcium uptake: possible mechanisms of control by stanniocalcin. Fish Physiol. Biochem. 11, 205–215.

Vincent, F. and Duncton, M. A. (2011). TRPV4 Agonists and Antagonists. Curr. Top. Med. Chem. 11, 2216–2226.

Volberg, T., Geiger, B., Kartenbeck, J. and Franke, W. W. (1986). Changes in membrane-microfilament interaction in intercellular adherens junctions upon removal of extracellular Ca^2+^ ions. J. Cell Biol. 102, 1832–1842.

Watabe-Uchida, M., Uchida, N., Imamura, Y., Nagafuchi, A., Fujimoto, K., Uemura, T., Vermeulen, S., Van Roy, F., Adamson, E. D. and Takeichi, M. (1998). α-Catenin-vinculin interaction functions to organize the apical junctional complex in epithelial cells. J. Cell Biol. 142, 847–857.

Welch, M. W., Hamar, D. W. and Fettman, M. J. (1990). Method comparison for calcium determination by flame atomic absorption spectrophotometry in the presence of phosphate. Clin. Chem. 36, 351–354.

Weng, L., Dai, H., Zhan, Y., He, Y., Stepaniants, S. B. and Bassett, D. E. (2006). Rosetta error model for gene expression analysis. Bioinformatics 22, 1111–1121.

West, M. R., Ferguson, D. J. P., Hart, V. J., Sanjar, S. and Man, Y. (2002). Maintenance of the epithelial barrier in a bronchial epithelial cell line in dependent on functional E-cadherin local to the tight junctions. Cell Commun. Adhes. 9, 29–44.

Wongdee, K. and Charoenphandhu, N. (2013). Regulation of epithelial calcium transport by prolactin: From fish to mammals. Gen. Comp. Endocrinol. 181, 235–240.

Yu, A. S. L., Cheng, M. H. and Coalson, R. D. (2010). Calcium inhibits paracellular sodium conductance through claudin-2 by competitive binding. J. Biol. Chem. 285, 37060–37069.

Zhang, Y., Sivasankar, S., Nelson, W. J. and Chu, S. (2009). Resolving cadherin interactions and binding cooperativity at the single-molecule level. Proc. Natl. Acad. Sci. U. S. A. 106, 109–114.

